# Pseudogene co-expression networks reveal a robust prognostic signature of survival in pediatric B-ALL

**DOI:** 10.1101/2025.03.14.643224

**Authors:** Arturo Kenzuke Nakamura-García, Marieke L. Kuijjer, Jesús Espinal-Enríquez

## Abstract

Risk classification in B-cell acute lymphoblastic leukemia (B-ALL) remains challenging, even in the era of genomic precision medicine. Current molecular classifiers fail to fully explain the heterogeneity in patient outcomes, suggesting that key regulatory layers remain hidden. Here, we uncover a previously unexplored dimension of B-ALL biology by analyzing co-expression patterns between pseudogenes using single-sample co-expression networks (n = 1,416). Principal component analysis showed that these interactions explain a major component of variability among patients and contribute to patient stratification into clusters with distinct overall survival. After identifying interactions associated with these clusters, we used a LASSO-based feature selection pipeline to derive a three-interaction signature that predicted patient survival, with *RPL7P10* –*RPS3AP36* emerging as the most robust biomarker. Our study shows that co-expression between pseudogenes represents a previously unrecognized layer of molecular heterogeneity in B-ALL, harboring promising molecular markers for future studies.

## Introduction

B-cell acute lymphoblastic leukemia (B-ALL) is a type of hematologic malignancy characterized by a clonal expansion of malignant B-cell progenitors. This type of leukemia is the most frequent among pediatric patients and, although it has high cure rates, relapse is still very common [1–3]. Next-generation sequencing has helped to uncover new B-ALL subtypes, which are mostly defined by gene fusions and chromosomal rearrangements[4–9]. However, these works have mainly focused on deregulation among protein-coding genes, leaving the potentially oncogenic roles of non-coding DNA sequences largely unexplored. Recent studies have started to explore the potential regulatory roles of pseudogenes, suggesting that they may contribute to the deregulation of the gene expression landscape observed in complex diseases such as B-ALL [10].

A pseudogene is a copy from a protein-coding gene that, due to detrimental alterations in its sequence, has lost its ability to code the original functional protein [11]. Historically, these type of DNA sequences have been considered “junk” DNA, and hence, research in their possible roles in gene regulation has been largely hampered. However, it has been demonstrated that pseudogenes can be transcribed and even translated, although their functions diverge from their parental coding gene [12]. Multiple works have demonstrated that pseudogenes are actively involved in gene regulatory circuits through diverse mechanisms. For example, they can alter the translation of their parental gene through endogenous competition for regulatory elements, or even alter DNA structure to promote or repress the transcription of a sequence (reviewed in [11]). Moreover, pseudogenes have been found to function as important modulators of gene expression in cancer [13]. Given the regulatory role of pseudogenes in gene expression, understanding their interactions within the co-expression landscape can provide valuable insights into the underlying biological processes in which they are involved. Furthermore, analyzing them in the context of complex diseases, like BALL, can reveal novel information about the regulatory re-wiring that drives disease pathology.

Gene co-expression networks (GCNs) are a common tool in the field of systems biology to approach the inherent complexity of biological systems[14]. These networks are theoretical graphs composed by genes connected between them when their expression profiles are correlated. A co-expression interaction between a pair of genes suggests a coordination which is possibly part of the same biological process [15]. Previous works have described GCNs from multiple cancer types[16–26].

We have reported an increased co-expression between pseudogenes in hematological cancers compared to normal samples [27]. This suggests that a coordination between these sequences could be involved in the biology of these malignancies, which warrants further exploration. However, these recent results were obtained through the analysis of aggregate gene co-expression networks, constructed by integrating data from multiple samples. Consequently, these pseudogene co-expression networks provide only a summary of the biological data across populations. As a result, the aggregate networks fail to capture the intrinsic biological heterogeneity within the population.

Biological samples are highly heterogeneous; understanding how their inherent variability affects biological systems is crucial for treatment design, or when evaluating patient clinical features (such as survival rates). In general, a one-size-fits-all approach is inadequate for addressing complex diseases [28, 29], necessitating a more personalized approach.

Because of the latter, in the context of systems biology, new approaches have been developed to construct gene co-expression networks at the single-sample resolution. These approaches enable the creation of “single-sample networks” (SSNs), which describe the coordination of genes within individual samples from a population. Various methods have been proposed for this purpose (reviewed in [30]), each differing in how the individual sample networks are calculated. One such method is LIONESS [31], which employs a leave-one-out strategy to estimate each sample’s contribution to the aggregate network. This method has been successfully applied to understand gene regulatory mechanisms in various cancers, leading to the identification of potential new regulatory subtypes in leiomyosarcoma [32], sex-biased regulatory patterns affecting prognosis in colon cancer [33], regulatory signatures associated with poor prognosis in glioblastoma [34], and subtype-specific coexpression hotspots in breast cancer [35].

Motivated by our previous observation of increased pseudogene co-expression in leukemias [27], we hypothesized that modeling these interactions at the single-sample level could reveal heterogeneity associated with clinical outcomes in B-ALL. To test this, we applied the LIONESS algorithm to construct single-sample co-expression networks from RNA-seq data of 132 patients from the TARGET-ALL-P2 cohort, focusing exclusively on pseudogene–pseudogene interactions. Based on edge weights from these individualized networks, we clustered patients, finding that pseudogene coordination patterns alone were sufficient to stratify patients into groups with significantly different overall survival (OS). Importantly, this signal was not recoverable when pseudogene interactions were excluded from the analysis.

We prioritized the most prognostically informative edges through a resampling-based LASSO strategy applied to the combined dataset of two cohorts (TARGET-P2-ALL and MP2PRT-ALL), which consistently selected three stable interactions. A multivariate Cox model built on these interactions, trained exclusively in TARGET and evaluated in MP2PRT, retained predictive performance and achieved significant OS stratification across cohorts. It is important to mention that MP2PRT-ALL cohort is enriched for patients with standard risk and high-risk with genetic features associated with favorable outcome, with only 6% mortality within five years.

To test whether the observed generalizability of the model was driven by a true biological signal or was merely an artifact of the feature selection pipeline, we generated 1,000 null models with randomly selected features, applying the same LASSO-based filtering and evaluation procedures. Only one of these null model outperformed the real model, highlighting the biological relevance of the selected pseudogene interactions.

Among the selected features, the interaction between *RPL7P10* and *RPS3AP36* stood out as the most consistent and predictive. We evaluated its prognostic value independently and showed that it was sufficient to significantly stratify patients by survival risk in both datasets.

Overall, these results highlight pseudogene co-expression as a potential source of novel survival biomarkers in cancer, and underscore the value of sample-level network approaches for advancing personalized medicine in B-ALL.

## Results

### Pseudogene co-expression landscape reveals inter-chromosomal and family-driven coordination

To characterize the transcriptional coordination among pseudogenes in B-ALL, we constructed an aggregate co-expression network using RNA-seq data from 132 patients of TARGET-ALL-P2 cohort (see Methods). Edges with a correlation p-value below 10*^−^*^8^ were considered significant. The resulting network included 45,114 edges and 11,379 genes, of which 6,032 edges connected 865 pseudogenes. This pseudogene co-expression network, referred to hereafter as *PG_net_*, forms the foundation for all subsequent analyses (Fig. 1).

**Fig. 1.**
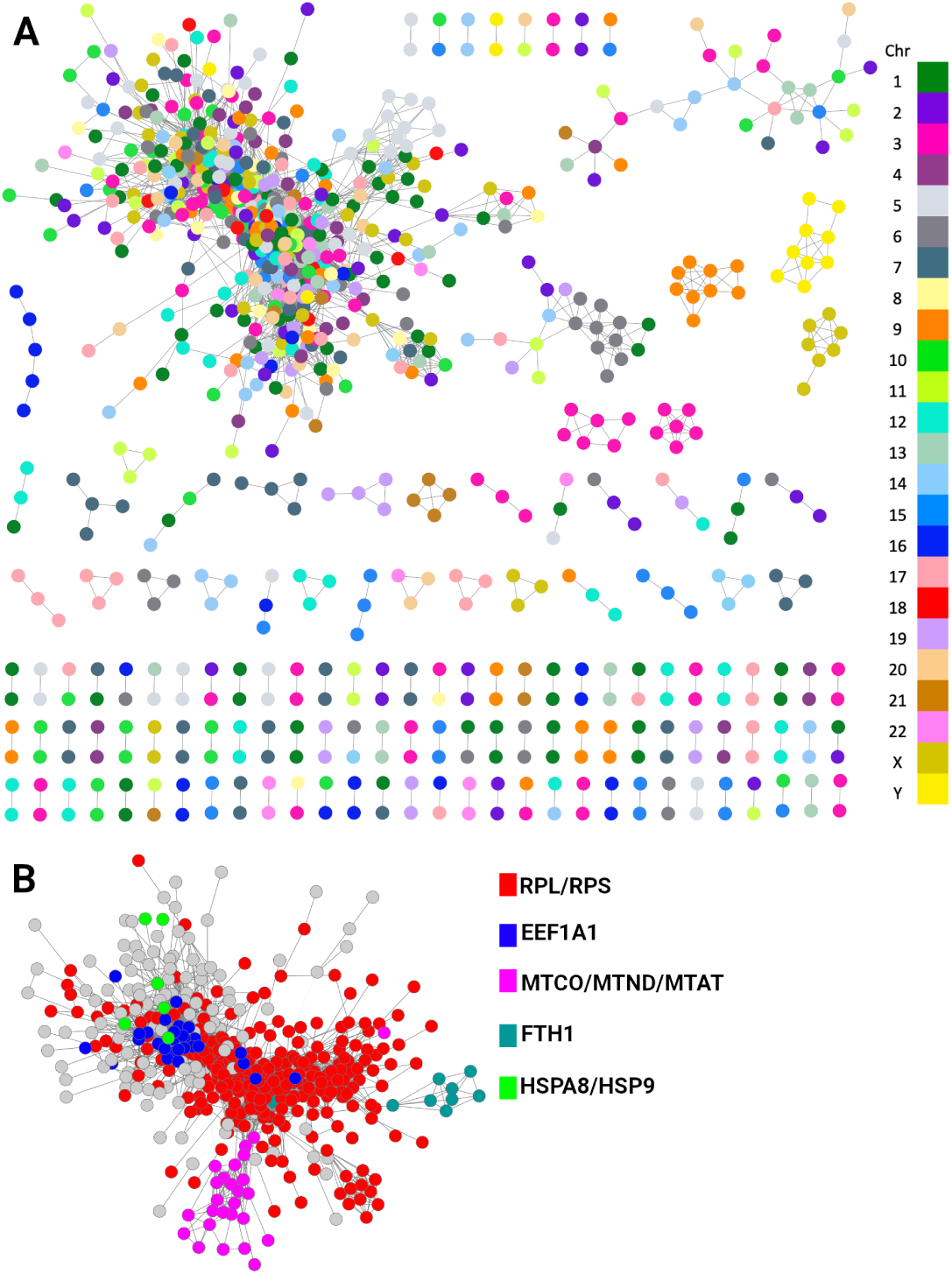
A) Aggregate gene co-expression network between pseudogenes (*PGnet*) in B-ALL samples. The color of nodes corresponds to the chromosome in which those pseudogenes are located. To note, the largest component contains pseudogenes from all chromosomes, meanwhile the smaller ones are mostly formed of pseudogenes from the same chromosome. B) Largest component of A), with pseudogenes colored according to their families.

In Fig. 1A, nodes correspond to pseudogenes and are colored according to their chromosome of origin. While smaller subgraphs are composed predominantly of intra-chromosomal edges, the largest component integrates pseudogenes from all chromosomes, suggesting broad inter-chromosomal coordination. However, when pseudogenes are colored by family (Fig. 1B), a clear clustering by parental origin emerges, with 94.6% of interactions in the largest component occurring between pseudogenes from different chromosomes but belonging to the same or closely related gene families. Given the sequence similarity between pseudogenes and their parental genes [36], we evaluated whether the observed network structure could be explained by misalignment artifacts in RNA-seq. To test this, we compared expression correlations and sequence similarity across pseudogene families (Fig. S1A). Fourteen families showed moderate associations (Fig. S1B–D); however, highly similar pseudogenes were frequently expressed at low levels (TPM ¡ 1; Fig. S1E). When these low-expression members were excluded, correlations weakened substantially and family sizes decreased (Fig. S1F), indicating that alignment artifacts are unlikely to account for the global coordination observed in *PG_net_*. Moreover, co-expression edge weights were essentially independent of sequence similarity between pseudogene pairs (Fig. S2), further supporting the biological relevance of the network.

### Individualized *P G_nets_* stratify B-ALL patients into survival-relevant clusters

To explore inter-patient heterogeneity in pseudogene coordination, we applied the LIONESS algorithm [31] to construct single-sample networks (SSNs) from the *PG_net_* for each of the 132 patients in the TARGET cohort. This approach estimates the contribution of each individual sample to the global co-expression network by calculating the difference in correlation values when the sample is excluded. The resulting SSNs allow the identification of patient-specific regulatory patterns, enabling fine-grained comparisons between individuals.

To identify subgroups of patients with similar pseudogene co-expression profiles, we used the M3C clustering algorithm, which estimates a Relative Clustering Stability Index (RCSI) and evaluates significance against a reference null distribution [37]. Since M3C recommends to use a set of variable features that are approximately normally distributed, we first filtered the top 25% most variable edges from the *PG_net_* and then scaled these edges to Z-scores across samples (by subtracting the mean edge weight from the sample weight and dividing it by the edge’s standard deviation).

This analysis identified three clusters (K = 3; Fig. 2A), which were distinguished by overall edge weight patterns: one with predominantly negative values (C1), a second with near-zero values (C2), and a third with highly positive weights (C3) (Fig. 2B). Strikingly, Kaplan-Meier analysis revealed that patients in C1 exhibited significantly better overall survival (OS) compared to those in C2 and C3 (Fig. 2C). These findings suggest that the coordination state of pseudogenes may be associated with differential prognosis.

**Fig. 2.**
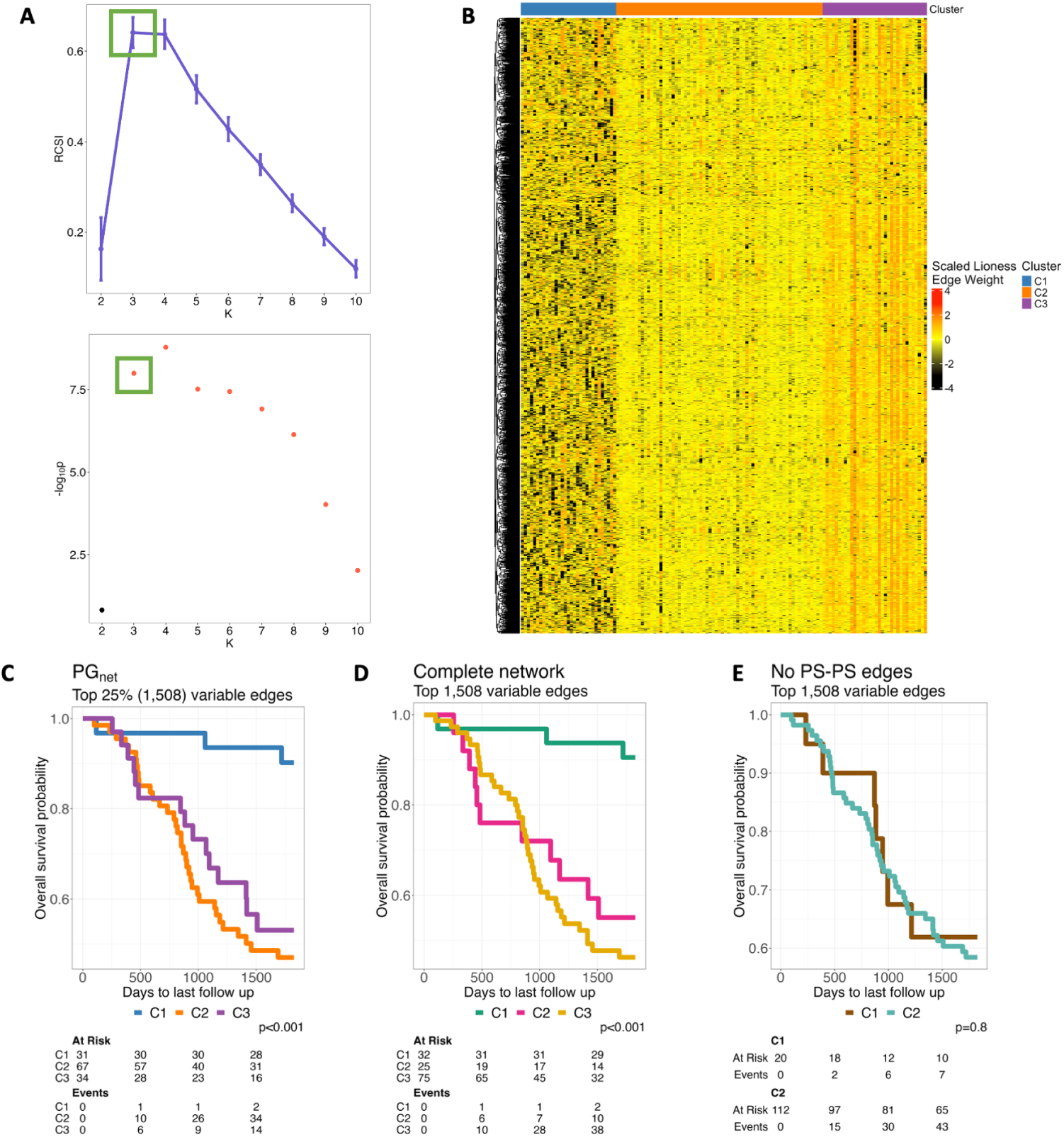
Clustering analysis of single sample networks of B-ALL samples from TARGET-ALL-P2. A) The M3C algorithm shows K = 3 as the optimal partition for the data. The top figure shows the RCSI scores calculated for up to 10 partitions of the dataset. The bottom figure shows the p-values associated with each K of the top figure. B) Heatmap showing the SSNs’ values scaled across samples clustered according to the M3C result. C) Kaplan-Meier analysis in the *PGnet*. D) KM plot in the network containing all types of edges. E) KM in the network without interactions between pseudogenes.

### Pseudogene-based network stratification is robust and reproducible across datasets

To ensure that the clustering patterns observed in *PG_net_* were not artifacts of network construction or statistical noise, we implemented a series of rigorous validation steps. First, we considered whether applying LIONESS to highly variable edges could artificially inflate sample-to-sample variability and, in turn, drive clustering independent of biological signal. To address this, we repeated the analysis using two alternative network versions: (1) a complete network including all genes and their significant interactions, and (2) a network excluding pseudogene-pseudogene (PS–PS) edges. Both networks were filtered to retain the top 1,508 most variable edges, matching the number used in the original *PG_net_* analysis. Clustering based on the complete network closely recapitulated the structure and survival separation observed in the *PG_net_* (Fig. 2D). In contrast, removing PS–PS edges resulted in weaker sample stratification, and the M3C algorithm identified only two clusters with no significant difference in survival (Fig. 2E). Principal component analysis confirmed that networks lacking PS–PS edges exhibited reduced inter-sample variance (Fig. S3), suggesting that pseudogene interactions capture a distinct layer of biological variability.

To formally evaluate whether clustering based on highly variable edges alone could lead to spurious survival associations, we generated 1,000 null models by randomly selecting 1,508 edges from the complete network. None of these models produced clusters with significant survival differences (Fig. S4), reinforcing that pseudogenedriven stratification reflects a non-random and biologically relevant signal.

We next tested the reproducibility of our findings in an independent dataset of 1,284 B-ALL samples from the MP2PRT-ALL project [38]. In this dataset, the *PG_net_*consisted of 19,244 edges among 1,763 pseudogenes. Clustering of the top 25% most variable edges from this *PG_net_* identified two patient groups with significantly different survival outcomes (Fig. S5A). Interestingly, networks excluding PS–PS edges also yielded survival-related clusters in this dataset (Fig. S5B–C), indicating that pseudogene interactions are not the sole contributors to prognostic stratification in this cohort.

### Differential co-expression analysis identifies 42 interactions associated with survival

To identify pseudogene–pseudogene edges that differentiate clusters based on survival, we performed a Wilcoxon rank-sum test on the scaled co-expression values across single-sample networks, comparing each edge between clusters. We also computed the median difference in edge weights between groups (e.g., median of C2 minus median of C1, *C*2 *− C*1).

We observed that the clusters associated with poorer survival exhibited overall higher co-expression scores (i.e., more positive edge weights, Fig. 3A,B and Fig.S6). Notably, in the TARGET dataset, the C2 vs. C1 comparison revealed both positively and negatively differential co-expression edges (Fig.3A,B), indicating heterogeneity in the dysregulation patterns.

**Fig. 3.**
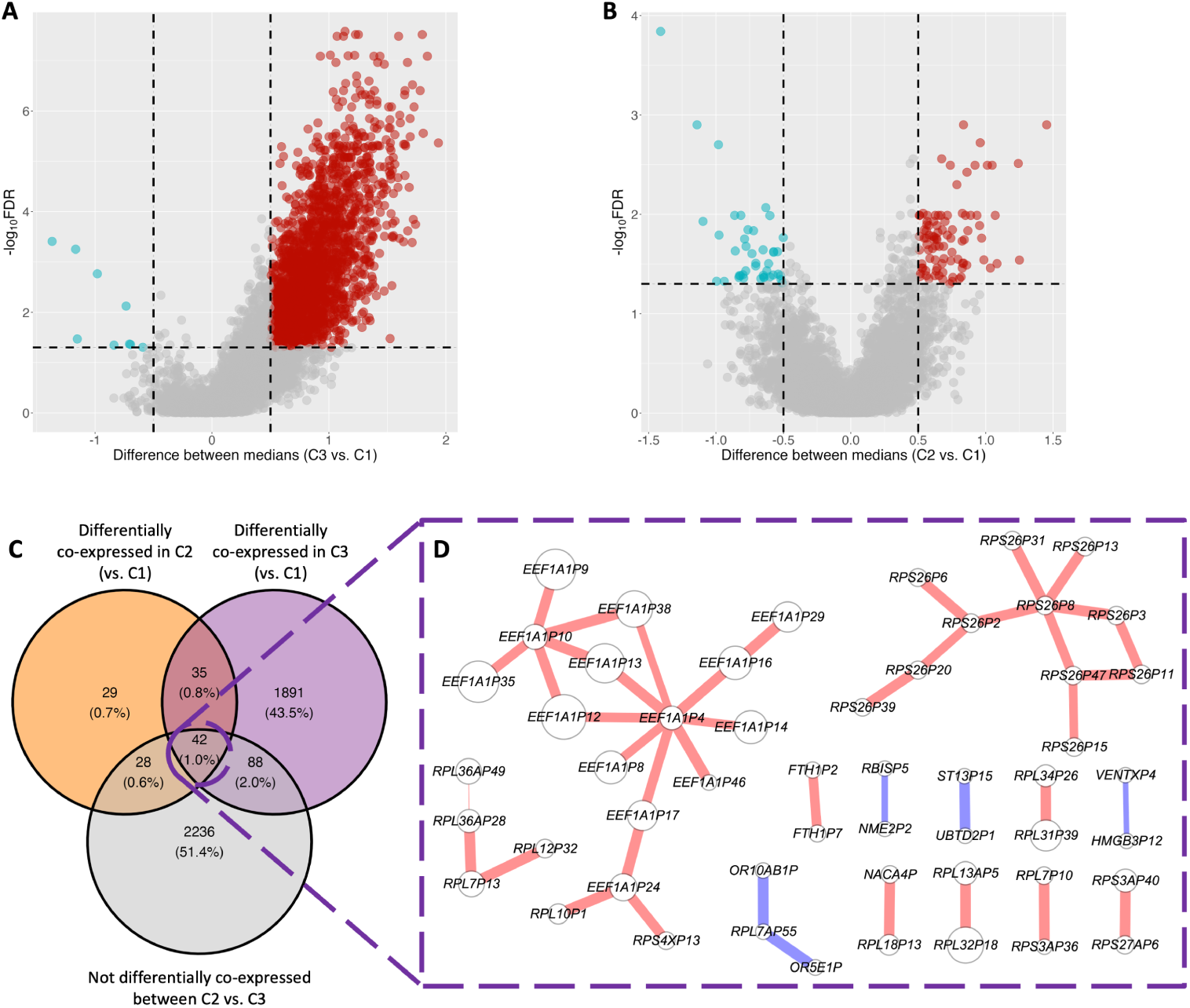
Volcano plots showing the differential co-expression analysis between Clusters from the *PGnets*, A) the comparison between C3 and C1. B) Comparison between C2 and C1. C) Venn diagram of differentially co-expressed edges between high (C1) and low survival (C2 and C3, orange and purple circles) clusters, versus not differentially co-expressed edges between C2 and C3. D) Resulting network from the intersection in C. Color of pseudogenes refers to their differential expression values between C3 V. C1: edge color shows the differential co-expression between edges of Clusters 3 V. 1. Size of nodes is proportional to their degree in the *PGnet*.

To focus on edges that robustly distinguish survival outcomes, we applied the following filtering strategy on the TARGET *PG_net_*:

- Edges had to show a significant difference in both the C3 vs. C1 and C2 vs. C1 comparisons.
- Edges had to show no significant difference between C3 and C2, as these two clusters

had similar overall survival and differences between them are unlikely to reflect survival-associated mechanisms (Fig. 3C).

This filtering resulted in a network of 42 pseudogene–pseudogene interactions (Fig.3D), which we selected for downstream analysis.

In the MP2PRT dataset, only two clusters were identified. Therefore, the differential co-expression analysis was limited to the C2 vs. C1 comparison (Fig.S6), which yielded a large number of significant edges. However, given the reduced heterogeneity in MP2PRT and the lower interpretability of its cluster-specific co-expression patterns, we focused subsequent analyses on the 42 pseudogene interactions identified in TARGET.

### Development of a pseudogene-based predictive model

So far, our analyses have shown that co-expression between pseudogenes is a widespread phenomenon in B-ALL and is significantly associated with patients’ overall survival. However, translating co-expression biomarkers into clinical practice remains challenging, as clinical risk stratification typically relies on variables of diverse nature, most often derived from cytogenetic or clinical assessments. A key limitation is that co-expression measurements, particularly those derived from LIONESS, are computed at the population level and are only interpretable within the context of the reference cohort used to build the network.

To obtain sample-specific co-expression values in the MP2PRT cohort that are comparable to those in TARGET, we projected each MP2PRT sample into the TARGET co-expression space (details in Methods). This allowed us to apply the previously 42 identified pseudogene interactions to MP2PRT in a consistent manner comparable to TARGET.

To select robust co-expression interactions for survival prediction, we combined the TARGET cohort with the projected MP2PRT samples and performed 100 rounds of LASSO-based feature selection by randomly selecting 50% of the samples.

Next, to select the most robust and minimal set of features for the multivariate model, we ranked all candidate edges by their selection frequency across the 100 LASSO partitions. From these iterations, we identified 3 with high selection stability across resampling and stable coefficient values (Fig. 4A; Fig. S7A). Specifically, we selected the interactions *RPL7P10* –*RPS3AP36*, *RPL36AP28* –*RPL7P13*, and *RPS26P2* –*RPS26P6* (details in Methods). This strategy allowed us to identify a consistent and minimal feature set under increased sample diversity, simulating a more realistic context where expression profiles may vary across patients or cohorts. To evaluate whether the predictive signal captured by the selected pseudo-gene interactions generalized beyond this combined cohort (TARGET + projected MP2PRT), we trained a Cox model exclusively in TARGET (Table 1) and tested its performance in the MP2PRT data using projected LIONESS edge weights. Risk scores were computed for all samples using the trained model, and patients were classified into lowand high-risk groups based on the median risk score derived from the TARGET cohort, which served as a fixed decision threshold across cohorts. This model consistently achieved significant separation of survival curves between groups across cohorts (Fig.5A and B), confirming that the model retains predictive value beyond the original training set (TARGET).

**Fig. 4.**
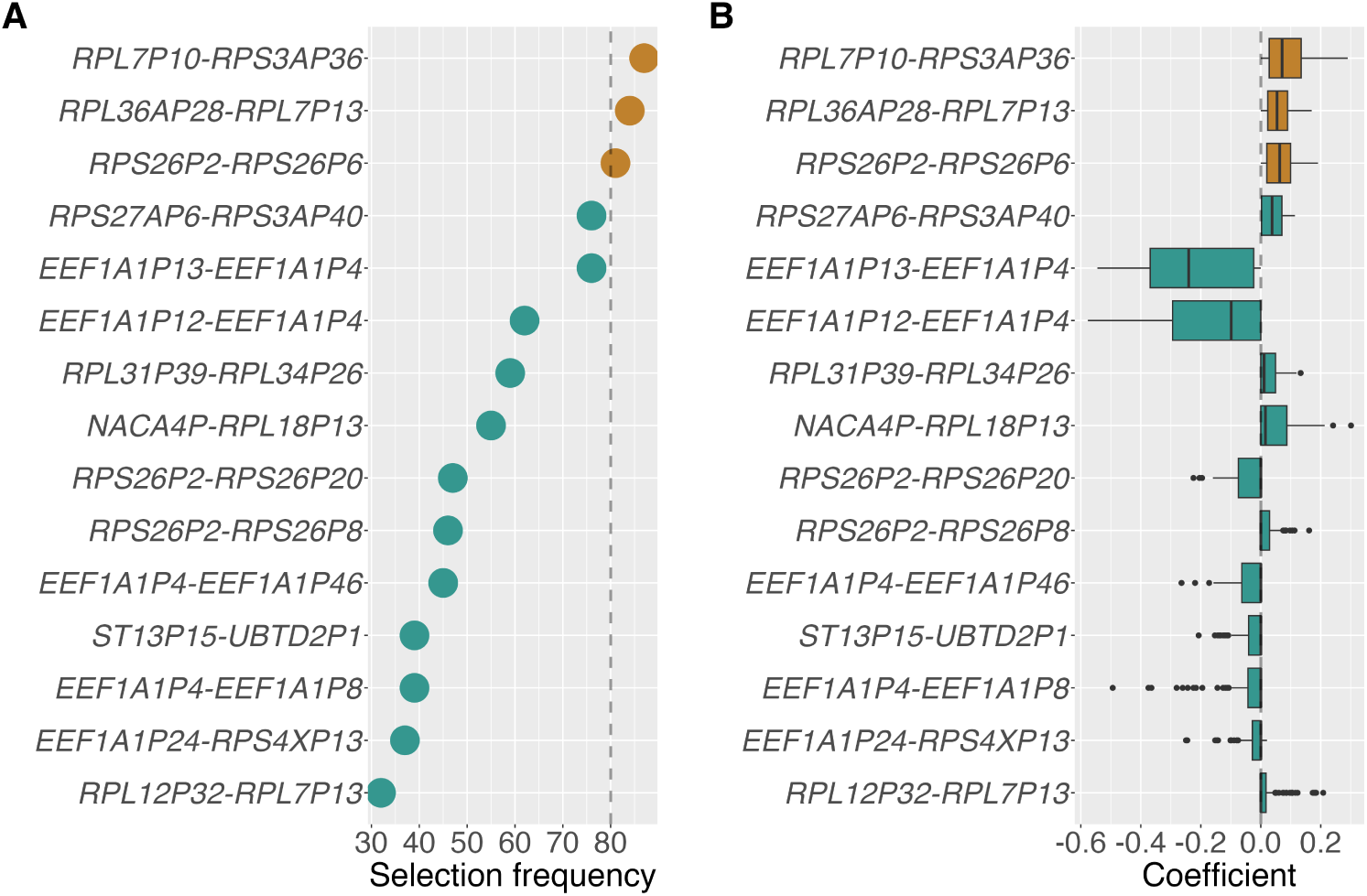
Feature selection frequency and model coefficients for pseudogene–pseudogene co-expression links. A) Frequency with which each edge was selected across 100 LASSO models trained in the discovery set. The interactions selected for downstream analysis are colored in in brown. B) Distribution of coefficients assigned to each edge across the 100 LASSO iterations. Colored boxes indicate the three final selected features. Both panels shows the top 15 interactions most frequently selected.

**Fig. 5.**
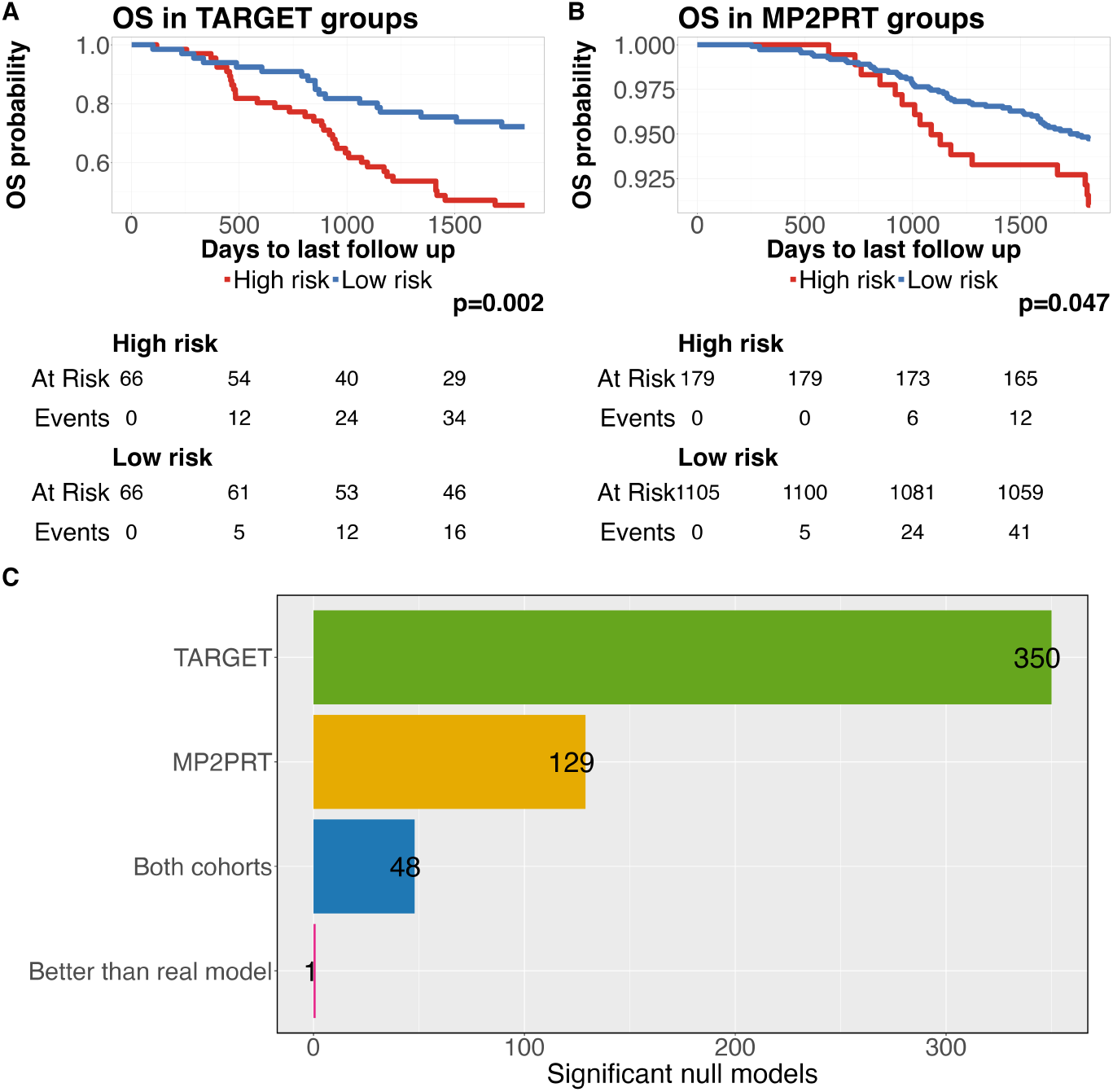
Predictive performance and validation of the pseudogene-based survival model. A) Kaplan–Meier curve in the TARGET cohort, stratified by the multivariate model. B) Equivalent analysis in the MP2PRT cohort, using the same stratification threshold defined in TARGET. C) Amount of significant null models across different evaluation cohorts and criteria.

**Table 1.** Cox proportional hazards model trained in the TARGET dataset using three pseudogene-pseudogene interactions.

| Edge | Coefficient ($\beta$) | HR | 95% CI | p-value |
| --- | --- | --- | --- | --- |
| *RPL7P10 – RPS3AP36* | **0.3992** | **1.49** | **1.01 – 2.19** | **0.045** |
| RPL7P13 – RPL36AP28 | 0.2848 | 1.33 | 0.95 – 1.88 | 0.099 |
| RPS26P2 – RPS26P6 | 0.2100 | 1.23 | 0.89 – 1.70 | 0.214 |

*Model fit metrics:* Concordance index = 0.645 Likelihood ratio test = 15.62 on 3 df, p = 0.001 Wald test = 15.05 on 3 df, p = 0.002 Score (logrank) test = 15.63 on 3 df, p = 0.001

Next, to evaluate whether the observed cross-cohort generalization of our model could be explained by the feature selection procedure, we generated 1,000 null models by randomly selecting 42 co-expression edges from the global TARGET network (see Methods). Each null model was subjected to the same LASSO-based feature selection process, followed by model training exclusively in TARGET and evaluation in MP2PRT.

Among these 1,000 null models, only 4.8% achieved generalization from TARGET into MP2PRT (Fig. 5C), underscoring how rarely random feature sets yield consistent performance across cohorts, even under combined-cohort selection. Importantly, only a single null model outperformed the real model in predictive performance (Fig. S8).

### *RPL7P10* –*RPS3AP36* exhibits consistent predictive value across multiple time points

To better understand the individual contribution of each component of the three interactions model, we performed a univariate evaluation of their predictive performance over time. We calculated time-dependent ROC-AUCs for 1-to 5-year overall survival in both the TARGET and MP2PRT (projected edge weights) cohorts. The results are summarized in Figure 6A.

**Fig. 6.**
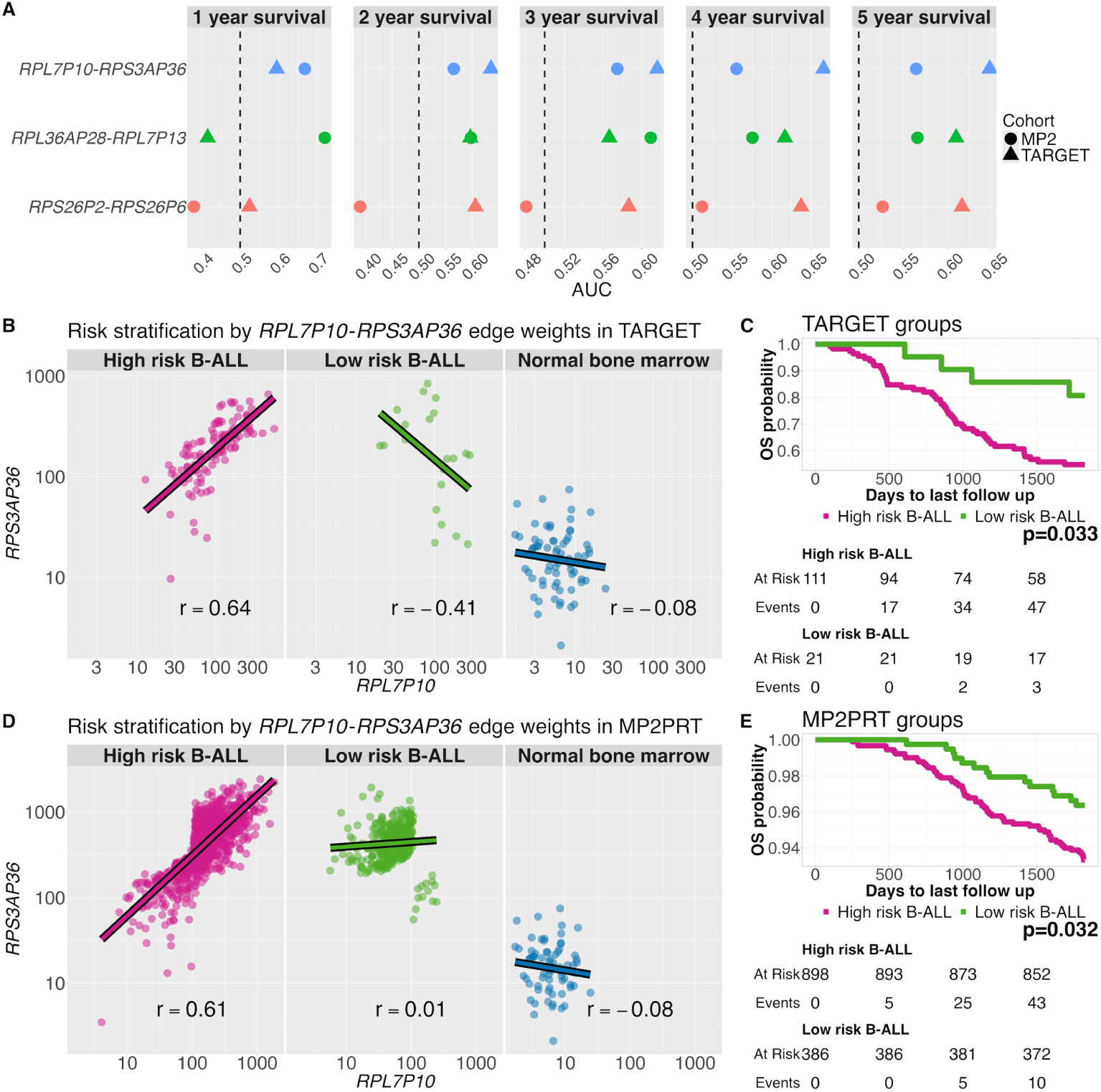
*RPL7P10* –*RPS3AP36* exhibits consistent prognostic value across cohorts. A) Timedependent ROC-AUCs for the three most frequently selected pseudogene interactions in both TARGET (triangles) and MP2PRT (circles) cohorts. *RPL7P10* –*RPS3AP36* consistently maintains *AUC >* 0.5 at all time points. B–C) Risk stratification in TARGET based on the edge weight of *RPL7P10* –*RPS3AP36*. B: expression relationship between *RPL7P10* and *RPS3AP36* in high-risk, low-risk, and normal bone marrow groups. C: the corresponding Kaplan–Meier curve for overall survival. D–E) Equivalent analysis in MP2PRT using projected data into the TARGET co-expression space. The edge weight stratifies patients into high-and low-risk groups with significant survival differences, recapitulating the patterns observed in TARGET.

Among the three pseudogene co-expression interactions, *RPL7P10* –*RPS3AP36* consistently achieved AUC values above 0.5 across all time points and in both cohorts, indicating a stable association with patient survival. In contrast, the other two interactions showed greater variability and lower AUC values, particularly in the MP2PRT cohort.

This consistent predictive behavior of *RPL7P10* –*RPS3AP36* over multiple time horizons and across datasets supports its potential as a robust individual biomarker.

Based on this observation, we selected this edge for further analysis to evaluate its prognostic value in greater detail.

To assess whether *RPL7P10* –*RPS3AP36* could independently stratify patients into prognostic groups, we classified samples based on the edge weight of this interaction. As a threshold, we used the median edge weight observed in the good prognosis group from the TARGET analysis, under the hypothesis that this value reflects a coordination state associated with favorable clinical outcomes. Using this fixed threshold, we stratified patients from TARGET and MP2PRT (using its projected edge weights) into”high-risk” (above the threshold) and”low-risk” (below the threshold) groups.

This strategy resulted in two clearly separable patient groups in both cohorts. In TARGET, the high-risk group showed a strong positive correlation between *RPL7P10* and *RPS3AP36* expression, while the low-risk group showed a weakly negative correlation. In MP2PRT, the high-risk group similarly exhibited strong positive correlation, whereas the low-risk group showed only weak coordination. Notably, in normal bone marrow samples, this edge showed a weak negative correlation, consistent with the pattern observed in the low-risk groups (Figure 6 B-E and Fig. S9).

## Discussion

To our knowledge, this is the first study to classify B-ALL patients solely based on co-expression interactions.

Pseudogene–pseudogene interactions remain poorly understood. Most prior research has focused on the interactions between pseudogenes with protein-coding genes or regulatory elements. For example, Carron et al. [39] explored pseudogene function through co-expression with coding genes, and Chang et al. [40] described ceRNA networks involving pseudogenes, lncRNAs, and miRNAs in lung cancer. These examples illustrate the emerging regulatory relevance of pseudogenes but leave a gap in understanding their internal coordination. Given their historical designation as “junk DNA,” we aimed to explore their potential as independent co-expression markers in leukemia.

Using data from 1,416 B-ALL patients from two cohorts (TARGET-ALL-P2 and MP2PRT-ALL), we constructed co-expression networks and single-sample networks (SSNs) focused on pseudogene–pseudogene (PG-PG) interactions. Clustering these SSNs revealed patient subgroups with significantly different overall survival in both datasets, underscoring the biological relevance of pseudogene coordination.

Interestingly, pseudogene-based networks (*PG_nets_*) exhibited higher variability than networks built from other gene types, suggesting that pseudogene co-expression captures underlying molecular heterogeneity among patients. In the TARGET cohort, clustering without pseudogene edges failed to identify survival-related subgroups, whereas networks including pseudogene interactions did reveal such stratification. Although networks excluding the PG-PG interactions in the MP2PRT cohort did yield survival-associated clusters, clusters formed using pseudogene edges alone also retained predictive value, reinforcing their clinical relevance in different clinical contexts.

Differences between the results across datasets could be associated to their clinical composition. MP2PRT includes nearly ten times more patients and is enriched for standardor high-risk cases with genetic features associated with favorable outcome, resulting in only 6% five-year mortality. By contrast, 40% of patients in the TARGET dataset died within five years. This clinical imbalance may partly explain discrepancies in the strength of the associations observed. Nonetheless, pseudogene co-expression consistently captured prognostic variation in both cohorts.

Notably, despite the clinical differences between TARGET and MP2PRT, we found consistent results across the two datasets. For example, heterogeneity in both cohorts is greatly explained by the pseudogenes co-expression (Fig. S3), and, per our clustering analysis, this heterogeneity is linked with the patients’ overall survival.

Additionally, using LASSO-based feature selection across multiple random partitions, we identified a pseudogene-based, minimal and robust signature to predict a patient’s survival. Among the three interactions included in the model, the one between *RPL7P10* and *RPS3AP36* stood out for its statistical significance and stability across time points and cohorts. When stratifying patients based on this edge (using a threshold derived from TARGET), we were able to distinguish high-and low-risk groups in both datasets. High-risk patients showed strong positive correlation, while low-risk and normal bone marrow samples exhibited weak or negative coordination.

Although the feature selection was performed on a combined dataset of TARGET and the projected MP2PRT networks to increase variability, the final model was trained exclusively in TARGET and evaluated independently in MP2PRT. To further ensure that the model’s generalization performance was not an artifact of the selection pipeline, we generated 1,000 null models using the same training and evaluation scheme. This analysis showed that the patterns found in TARGET rarely generalize into the MP2PRT cohort (only 4.8% of models).

Overall, these results support the robustness and specificity of the multivariate model and demonstrate that the selection pipeline does not, by default, yield predictive models across cohorts. Although using data from two cohorts for feature selection could introduce information leakage into the final model, our results show that this is not sufficient to generate models with cross-cohort stability. Moreover, this indicates that the predictive power of our pseudogene-based model is mainly driven by a biological signal rather than by artifacts of the selection procedure, highlighting the utility of our strategy for selecting stable features across cohorts.

Beyond its contribution to the multivariate model, the edge *RPL7P10* –*RPS3AP36* also demonstrated independent prognostic value when used as a single-feature classifier. Using the median edge weight observed in the low-risk TARGET group (C1 in Fig. 2C) as a threshold, hypothesized to reflect a favorable coordination state, we stratified patients in both cohorts into high- and low-risk groups. This threshold yielded clearly separable groups in both datasets, with significant survival differences. High-risk groups exhibited strong positive correlation between *RPL7P10* and *RPS3AP36*, whereas low-risk groups showed weak (in MP2PRT) and mildly negative coordination (in TARGET). Interestingly, normal bone marrow samples also showed negative correlation, mirroring the low-risk profile. These findings suggest that positive coordination between this pair of pseudogenes is a leukemia-specific feature associated with adverse prognosis, while loss or reversal of coordination may indicate a physiologic or less aggressive state.

Importantly, our findings suggest the existence of a coordination mechanism among pseudogenes that operates independently of their expression levels and may contribute to more aggressive tumor phenotypes. In our previous work on hematological malignancies [27], we observed that co-expression interactions predominantly occurred between pseudogenes from different chromosomes, in contrast to coding genes, which tended to interact within the same chromosome. This pattern suggests that the coordination of pseudogenes may be governed by distinct (and possibly opposing) mechanisms to those driving the loss of inter-chromosomal communication typically observed in cancer [19, 23, 41, 42].

A possible explanation for the co-expression (or even detected expression) of pseudogenes is a bias in RNA-seq read mapping due to high sequence similarity between pseudogenes and their parental genes. We addressed this possibility here and found no strong signs that our results were driven by this issue, supporting the biological relevance of the observed patterns.

Upon further testing, these co-expression biomarkers proposed in this work could become part of novel prognostic models. Our approach to analyze the risk of new samples relative to a reference dataset provides an example of how co-expression signatures can be used to classify new patients. Future research could focus on developing co-expression biomarkers using more accessible technologies than RNA-Seq, such as PCR, to facilitate their adoption in clinical settings. In addition, these new classification models could be integrated with drug repositioning approaches to recommend personalized therapies for high-risk patients, thus indirectly finding drugs that possibly target the co-expression biomarkers.

Further investigation involves exploring targeted therapies to modulate the highrisk coordination state of gene pairs. Identifying drivers of a high-risk coordination state, such as transcription factors (TFs) or other regulatory elements, could provide a means to alter these interactions.

Finally, integrating these perspectives into the field of network medicine could lead to novel strategies for risk stratification and therapeutic interventions. For instance, expanding traditional drug recommendation tools to incorporate co-expression and regulatory network biomarkers, would enable the identification of drugs capable of rewiring a patient’s co-expression network to improve treatment response and prognosis.

### Concluding remarks

This study reveals that pseudogene–pseudogene co-expression captures a previously unrecognized layer of prognostic information in B-ALL. By focusing on individualized networks derived exclusively from pseudogene interactions, we identified patient subgroups associated with distinct survival outcomes across independent cohorts. Importantly, we defined a minimal pseudogene co-expression signature that stratifies patients into highand low-risk groups and retains predictive value beyond its training cohort. Among the interactions in this signature, the one between *RPL7P10* and *RPS3AP36* emerged as a particularly robust biomarker. Together, our findings position pseudogene co-expression as a novel molecular feature with potential clinical utility for risk stratification and therapeutic development in B-ALL.

## Methods

The code for reproducing the results of this work can be found at: https://github. com/AKNG97/PSs-SSNs-in-B-ALL.git. Every step of the pipeline was executed in R.

## Data acquisition

The RNA-seq data used in this work were retrieved from Genomic Data Commons (GDC, https://portal.gdc.cancer.gov). Expression data from B-ALL samples were obtained from the MP2PRT-ALL and TARGET-ALL-P2 projects. Only cancer samples categorized as “Primary Blood-Derived Cancer - Bone Marrow” with complete survival follow-up information were included. Normal bone marrow samples were obtained from the TARGET-AML project. The types of counts used included TPM and unstranded counts.

A complementary gene annotation file was obtained from Biomart [43].

## Data pre-processing

Unstranded counts from cancer and normal samples were processed together. Only genes present in both the expression matrix and the external gene annotation were considered. All genes with duplicated HGNC symbols were removed. When multiple replicates of a sample were found, we summed the expression of genes between replicates with the function”collapseReplicates” from the”DESeq2” [44] package. Next, we filtered out lowly expressed genes in the following order:

1. Genes on the first quartile of mean expression.
2. Genes with 0 counts in at least 50% + 1 samples.
3. Genes with average expression less than 10.

### Network inference

For the co-expression analysis, raw counts were normalized with the following pipeline:

1. Within sample normalization by gene length and GC content with the EDASeq package [45].
2. Between sample normalization by trimmed mean of M values (TMM) with the NOISeq package [46].
3. Batch effect removal by ARSyN from the NOISeq package [46].

Gene lenght and GC content were retrieved from Biomart. Normalized counts from cancer samples were used to calculate the aggregate gene co-expression network using the Spearman’s rank correlation coefficient. Correlations with a p-value *<* 10*^−^*^8^ were considered as statistically significant. The LIONESS equation [31] was applied on the aggregate gene co-expression network to obtain the single-sample edge weights.

### Sequence similarity analysis in pseudogene families

The analysis between pseudogenes and parental genes used the TPMs counts for the TARGET B-ALL samples. Sequences were obtained from Biomart [43] using the function “getSequence” by Ensembl IDs. The type of sequence retrieved was “gene exon intron”. Sequence similarity scores were calculated using the Smith-Waterman algorithm with the function “smith waterman” from the “text.alignment” package [47].

For the analysis of sequence similarity between pairs of pseudogenes, we analyzed communities from the *PG_net_* identified through the Louvain algorithm (from the “igraph” [48] package) to generate the communities.

### Clustering analysis

Hierarchical clustering analysis was performed using the M3C package. The function uses a Monte Carlo simulation to generate a null distribution of stability scores while preserving the structure of the input data. From this, a Relative Clustering Stability Index (RCSI) is calculated to evaluate the plausibility of different numbers of partitions (K) in the data compared to the null hypothesis that the data contains no clusters (K=1) [37]. The function returns an optimal value of K using the RCSI and its associated p-value.

M3C was run with a set of top variable features normalized to Z-scores across samples. For the analysis of the *PG_net_* we used the top 25% most variable edges. Then, for the analyses of the complete network and the network without PG-PG edges, we used the same amount of top variable features as in the analysis of *PG_net_*. PCA analysis of input data is computed automatically in each run of M3C.

### Differential co-expression analysis

Differential co-expression analysis was performed by the median differences between groups. The significance of these differences was assessed using the Wilcoxon test. P-values were adjusted for multiple testing using the false discovery rate (FDR) method. Edges with an absolute median difference greater than 0.5 and an FDR *<* 0.05 were considered differentially co-expressed.

### Projection of the MP2PRT-ALL single-sample coexpression values into the TARGET-ALL-P2 LIONESS model

LIONESS models estimate the contribution of individual samples to the global coex-pression pattern by leveraging the variability present in a reference population. However, LIONESS-derived networks are not directly comparable when computed independently across different datasets.

Although one potential strategy for comparison would be to combine all samples into a single LIONESS model, this approach would be suboptimal in our context due to the distinct clinical composition of the TARGET and MP2PRT cohorts. Integrating both datasets into a unified model would introduce a substantial bias toward the coex-pression patterns of MP2PRT, given its larger sample size. Moreover, such integration would distort the interpretation of LIONESS edge weights derived from the original, independent models, complicating their biological interpretation and undermining the comparability of results across cohorts. For instance, it would not be possible to determine whether biomarkers identified in one cohort are also equally relevant in the other, as the combined model would alter the underlying edge weight distributions and their interpretation.

To overcome this limitation, we simulated an *in-silico* clinical setting by projecting each MP2PRT sample into the coexpression space defined by the TARGET dataset. Specifically, for each MP2PRT sample, we normalized its raw expression counts using the raw counts from TARGET, and then recalculated the LIONESS model incorporating that sample alongside the TARGET reference. This process was repeated independently for each MP2PRT sample to ensure the overall structure of the TARGET-based model remained stable. The resulting edge weights represent how each MP2PRT sample aligns with the TARGET coexpression landscape and are referred to as”projected” edge weights. By doing so, all samples are evaluated within a common reference framework, enabling a consistent comparison across cohorts while preserving the original interpretation of LIONESS edge weights.

### Feature selection and multivariate model construction

To build a minimal and robust predictive model for survival, we performed LASSO regression using the 42 edges that were differentially coexpressed between Cluster 1 (associated with good survival) and Clusters 2 and 3 (associated with poor survival) in the TARGET analysis.

To increase input variability, we included the projected MP2PRT edge weights alongside the original TARGET edge weights in a combined dataset. We then generated 100 random partitions, each including 50% of the total samples, and performed LASSO feature selection independently in each partition. This strategy served two complementary purposes.

First, randomly combining both datasets increased the overall variability in coexpression patterns, reducing the risk of overfitting to cohort-specific noise and enabling the identification of features informative across a broader range of biological and clinical heterogeneity.

Second, by incorporating MP2PRT samples into the selection process, we aimed to identify coexpression edges that were not only statistically robust within the TARGET cohort, but also stable under variation introduced by an independent population. In this sense, the combined selection process served as a filter to prioritize edges with greater potential to generalize beyond the original training cohort.

Finally, edges were ranked according to their selection frequency across the 100 LASSO runs. We then examined the change in frequency between consecutively ranked features to identify the first major inflection point in selection stability. This inflection occurred at the fourth-ranked position, where two edges tied with a selection frequency of 76. The drop from the first to the second and from the second to the third ranked edges was 3 in both cases, reflecting a stable decline. In contrast, the drop from the third to the fourth position was 5, representing a 1.67-fold increase in the magnitude of the frequency drop. Features ranked above this inflection point were retained for model construction, thereby minimizing the inclusion of unstable edges.

Although MP2PRT samples contributed to the feature selection process, the final multivariate Cox proportional hazards model was trained exclusively on TARGET samples, ensuring that model fitting remained independent from the evaluation cohort.

### Multivariate model performance evaluation

To evaluate the predictive power of the final model, we first used it to calculate risk scores for each sample in both the TARGET and MP2PRT cohorts. To define high- and low-risk groups, we applied a fixed threshold corresponding to the median risk score observed in the TARGET cohort. This approach ensured consistency in group assignment across datasets and avoided information leakage from the validation cohort. Additionally, using a fixed decision boundary allowed us to directly assess whether the survival-associated patterns identified in TARGET generalized to MP2PRT. We then computed time-dependent ROC-AUC values at 5 years to evaluate the model’s predictive accuracy in both cohorts.

Because the MP2PRT cohort was included in the feature selection process, we sought to determine whether the model’s performance in MP2PRT could be attributed to selection bias rather than to a truly generalizable biological signal. To address this, we constructed 1,000 null models by randomly selecting 42 edges from the full coexpression network defined in the TARGET dataset. For each null model, the projected edge weights for MP2PRT were computed, and the combined TARGET + MP2PRT dataset was subjected to the same LASSO-based feature selection procedure used in the real model, with 100 random partitions each including 50% of the samples.

Within each null model, edges were ranked by their selection frequency, and the drop in frequency between consecutively ranked edges was used to identify the first major inflection point in selection stability. Specifically, we computed the ratio between successive drops and retained all edges ranked above the point where the drop magnitude increased by at least 1.67-fold compared to the previous drop. This threshold was empirically derived from the behavior observed in the real model, where the top three features showed regular frequency drops of 3, followed by a sharper drop of 5 at the fourth position (5/3 = 1.67), marking the first disruption in selection stability. The same 1.67 ratio was applied across all null models to ensure consistency and comparability in feature selection.

A multivariate Cox proportional hazards model was then trained using only TARGET samples and the selected edges from each null model. Risk scores were calculated in both cohorts using the trained model, and samples were classified into highand low-risk groups using the same median threshold defined in TARGET. Each null model was evaluated using the Kaplan–Meier analysis, in which a model was considered significant if the analysis showed a p *<* 0.05 and the risk groups preserved the expected survival direction (i.e., higher survival in the low-risk group) across cohorts. Finally, to compare the overall performance of the null models against the pseudogene-based model we analyzed the models’ 5 year ROC-AUC values. Overall, this analysis showed that only 48 random models managed to generalize from TARGET into MP2PRT, and only one of them outperformed the real model by its 5 year ROC-AUC values.

This analysis enabled us to compare the real model’s performance to the distribution of null models under the same selection and evaluation criteria, and to determine whether the observed generalization in MP2PRT could be an artifact of the feature selection process.

### *RPL7P10* –*RPS3AP36* predictive analysis

To assess the individual contribution of the edges included in the multivariate model, we independently evaluated their predictive performance. Specifically, we computed time-dependent ROC-AUC values from years 1 to 5 for each edge. This analysis revealed that the interaction *RPL7P10* –*RPS3AP36* was the most stable and consistent predictor of overall survival across both cohorts.

To further evaluate the predictive value of this edge, we stratified the TARGET cohort into highand low-risk groups using its edge weights. Since this analysis was based on a single feature, we hypothesized that a threshold informed by underlying biological structure, rather than the median weight from the full distribution, would better distinguish the risk groups. For this purpose, we used the median edge weight observed within Cluster 1 (low-risk group) from the TARGET stratification. Based on our previous observations, we hypothesized that this value (–0.741) reflects a biologically relevant boundary that separates different coordination states associated with distinct clinical outcomes. This threshold was applied to both TARGET and MP2PRT (using projected edge weights) to define risk groups, which were then evaluated by Kaplan–Meier analysis for overall survival.

## Data Availability Statement

Clinical and RNA-sequencing (RNA-Seq) data corresponding to patients within the TARGET-ALL-P2, MP2PRT-ALL and TARGETAML projects in the Genomic Data Commons (GDC): https://portal.gdc.cancer.gov/. The pipeline for reproducing the results from this manuscript is available on GitHub and can be found at https://github.com/AKNG97/PSs-SSNs-in-B-ALL.git

## Acknowledgements

Arturo Kenzuke Nakamura-Garcia is a doctoral student from Programa de Doctorado en Ciencias Biomédicas, Universidad Nacional Autónoma de México (UNAM). This work is part of his PhD thesis. AKNG received a PhD scholarship from SECIHTI (number 806341). Financial support was received from SECIHTI (Grant number 237-2025), as well as Alianza Médica por la Salud A.C. (AMSA) 2025, to JEE.

## Supplementary information

- **Supplementary Table S1.** Spearman correlation between expression values and sequence similarity (between pseudogenes and parental genes) by family.
- **Supplementary Fig. S1.** Correlation analysis between sequence similarity and mean expression (TPMs) of pseudogenes with their parental genes in the TARGET dataset. A)Scatter plot of p-value and correlation value of analyzed families. Families of parental genes and pseudogenes in which a significant correlation (-log10(p-value) *<* 2 & absolute Rho *>* 0.5) was found are shown in blue. B), C) and D) show scatter plots of sequence similarity (between each PS and its parental gene) and mean expression, PSs above 1 TPM are shown in red and PSs below 1 in blue. E) Proportions of members with high sequence similarity (greater than 0.5) above and below detection level (1 TPM) from the 14 families with significant correlations (A). F) Spearman’s correlation coefficient when all the members are considered (full data) versus when only members above detection level are considered; here, only families from E) in which at least 10 members were above 1 TPM were considered.
- **Supplementary Fig. S2.** Correlation analysis between sequence similarity and edge weight in the aggregated network of the TARGET dataset. A to G show scatter plots by community. H) show the scatter plot of the complete network (6,032 edges).
- **Supplementary Fig. S3.** Principal components analysis performed on different network subsets. Top figures show the analysis on the TARGET network, bottom figures show the analysis using the MP2PRT data.
- **Supplementary Fig. S4.** Histogram of p-values from Kaplan-Meier analysis of clusters formed by randomly subsetting (1,000 times) 1,508 edges from the complete TARGET network.
- **Supplementary Fig. S5.** Kaplan-Meier analysis of clusters in MP2PRT data. A) Top 25% (4,811 edges) most variable PG-PG edges B) Top 4,811 most variable edges from the network without PG-PG edges. C) Top 4,811 most variable edges from the complete network containing all classes of edges.
- **Supplementary Fig. S6.** Volcano plot showing the differential co-expression analysis between clusters in the MP2PRT data formed by the analysis of the PGnets.
- **Supplementary Fig. S7.** Feature stability across random partitions. A) Features were ranked by how frequently they were selected across 100 LASSO- based feature selection iterations using 50%-sampled random partitions. B) First-order differences in selection frequency between consecutive features.
- **Supplementary Fig. S8.** 5-year ROC-AUC values in both TARGET and MP2PRT for the 48 significant null models. Dashed lines represent the performance of the real model.
- **Supplementary Fig. S9.** Coefficients of Pearson (A) and Spearman (B) correla-tions between samples of high and low survival risk by stratification using the single sample edge weights of *RPL7P10* -*RPS3AP36*.

