## supplementary figures for "Pseudogene co-expression networks reveal a robust prognostic signature of survival in pediatric B-ALL"

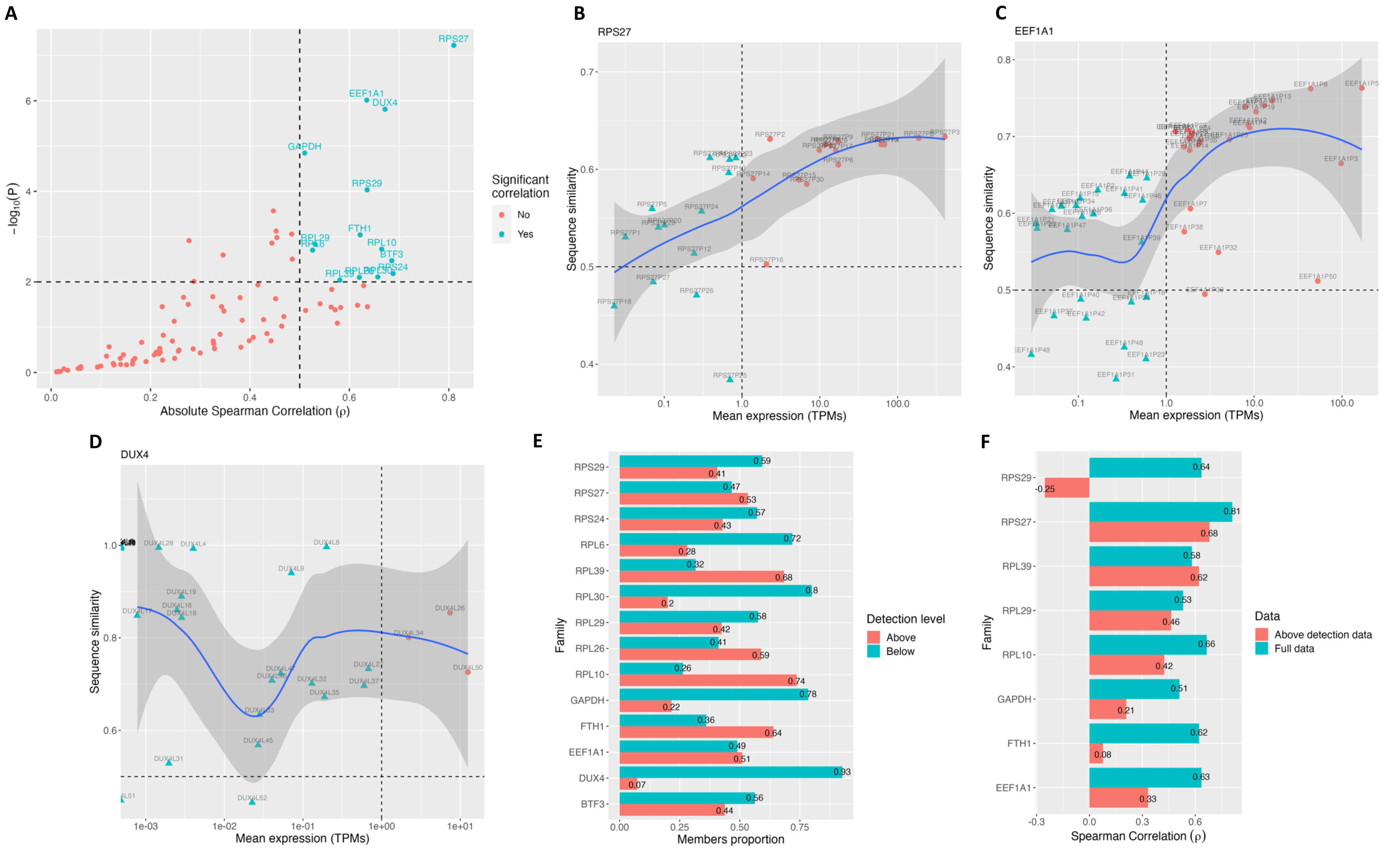


**Fig. S1.** Correlation analysis between sequence similarity and mean expression (TPMs) of pseudogenes with their parental genes in the TARGET dataset. **A)** Scatter plot of p-value and correlation value of analyzed families. Families of parental genes and pseudogenes in which a significant correlation (-log$_{10}$(p-value) < 2 & absolute Rho > 0.5) was found are shown in blue. **B)**, **C)** and **D)** show scatter plots of sequence similarity (between each PS and its parental gene) and mean expression, PSs above 1 TPM are shown in red and PSs below 1 in blue. **E)** Proportions of members with high sequence similarity (greater than 0.5) above and below detection level (1 TPM) from the 14 families with significant correlations (A). **F)** Spearman's correlation coefficient when all the members are considered (full data) versus when only members above detection level are considered; here, only families from E) in which at least 10 members were above 1 TPM were considered.

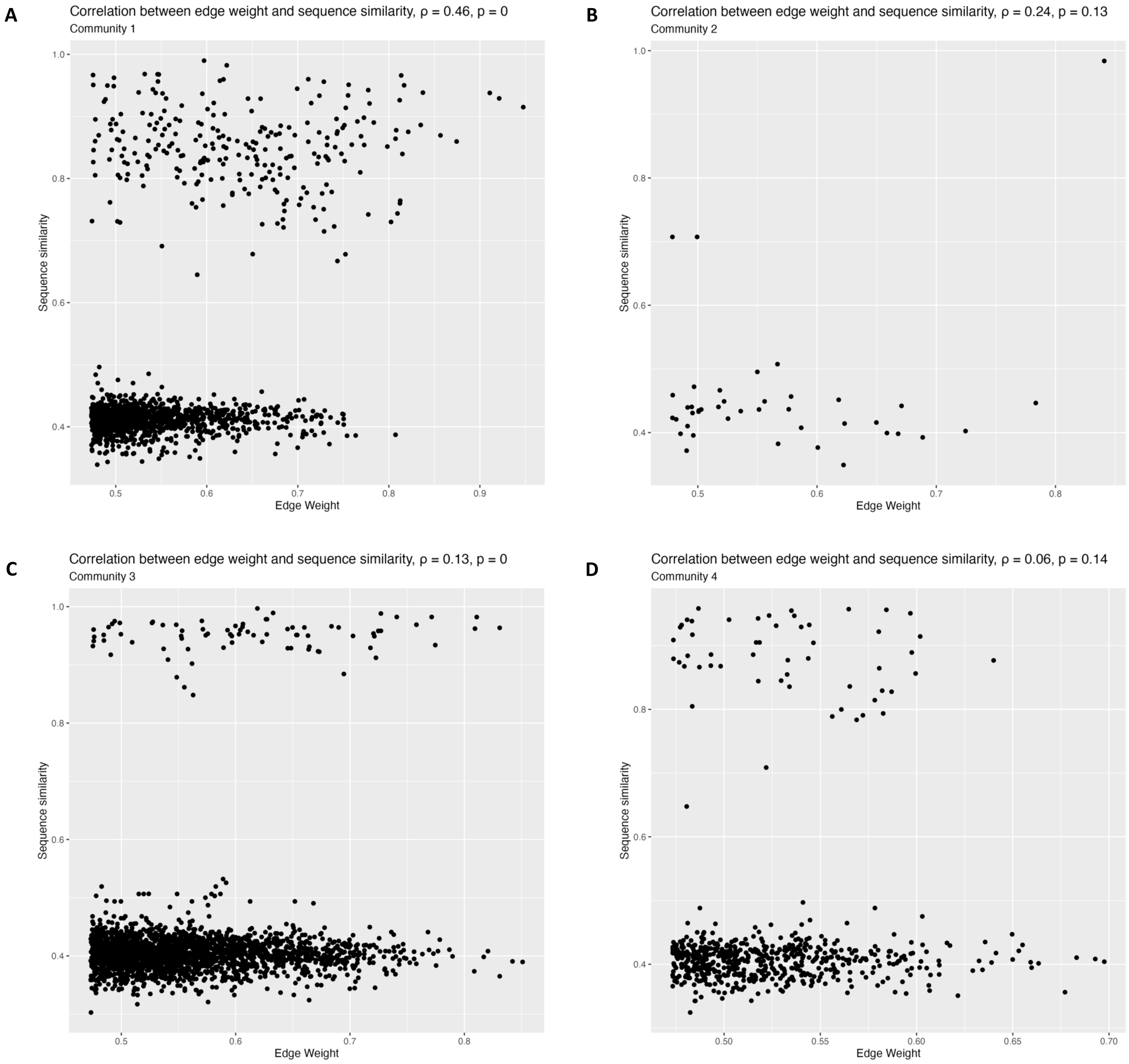
A
Correlation between edge weight and sequence similarity, ρ = 0.46, p = 0
Community 1
Sequence similarity
Edge Weight
B
Correlation between edge weight and sequence similarity, ρ = 0.24, p = 0.13
Community 2
Sequence similarity
Edge Weight
C
Correlation between edge weight and sequence similarity, ρ = 0.13, p = 0
Community 3
Sequence similarity
Edge Weight
D
Correlation between edge weight and sequence similarity, ρ = 0.06, p = 0.14
Community 4
Sequence similarity
Edge Weight

E

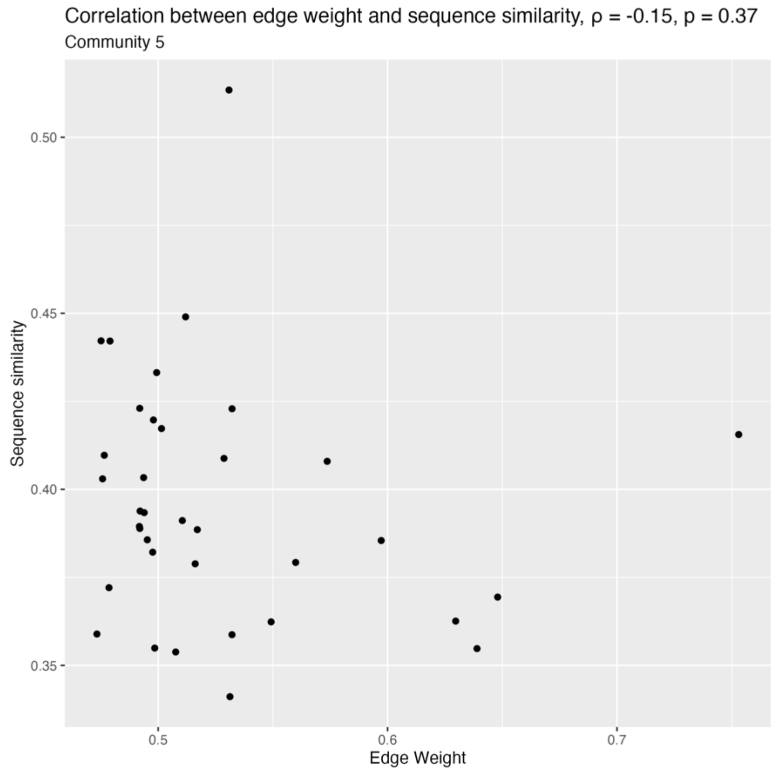
Correlation between edge weight and sequence similarity, ρ = -0.15, p = 0.37
Community 5
Sequence similarity
Edge Weight

F

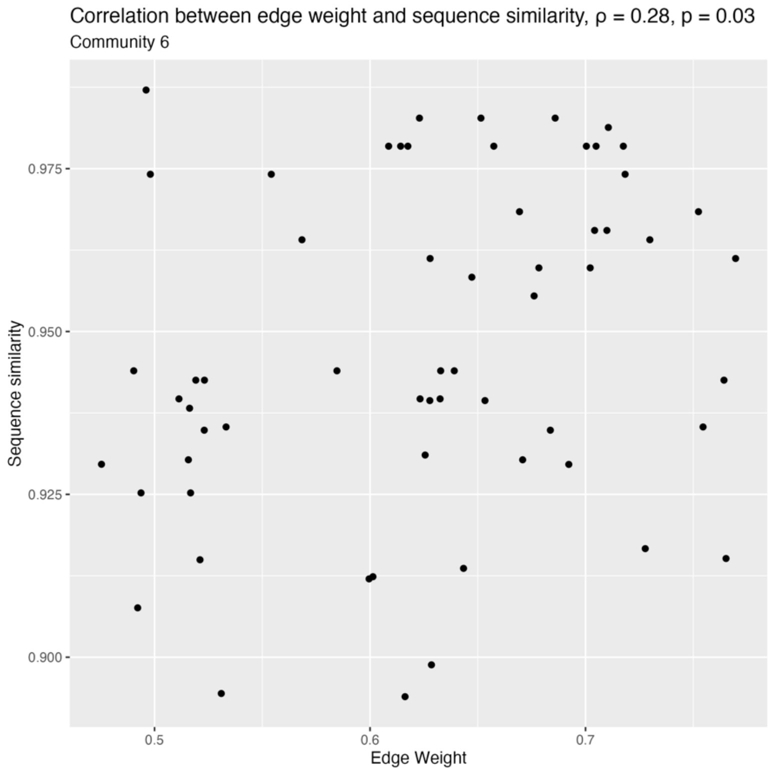
Correlation between edge weight and sequence similarity, ρ = 0.28, p = 0.03
Community 6
Sequence similarity
Edge Weight

G

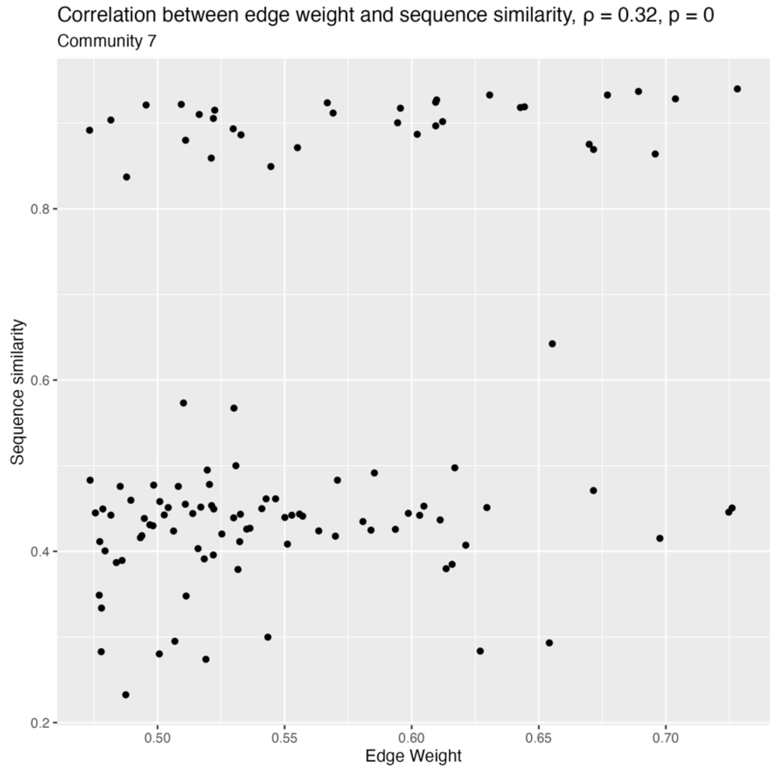
Correlation between edge weight and sequence similarity, ρ = 0.32, p = 0
Community 7
Sequence similarity
Edge Weight

H

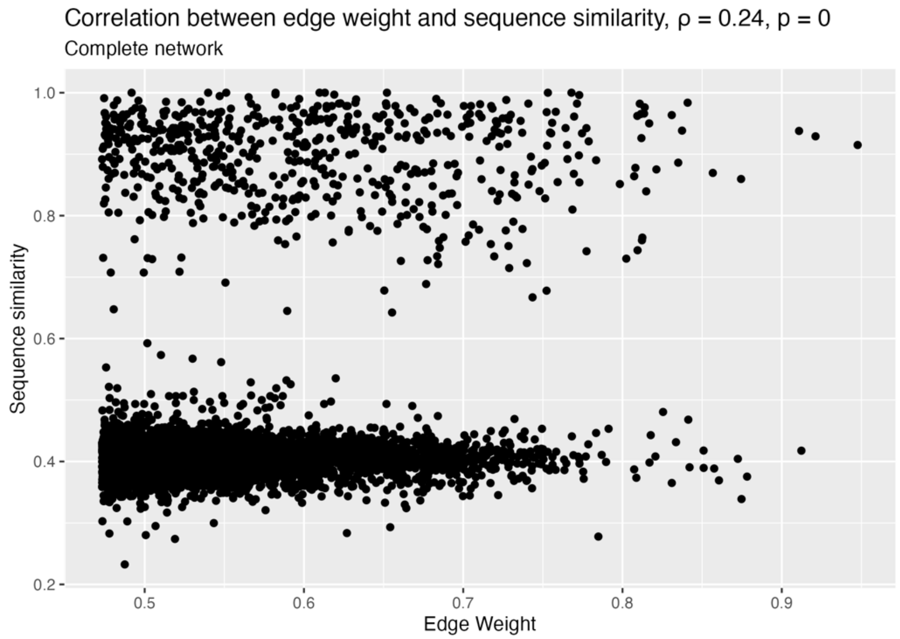
Correlation between edge weight and sequence similarity, ρ = 0.24, p = 0
Complete network
Sequence similarity
Edge Weight

**Fig. S2.** Correlation analysis between sequence similarity and edge weight in the aggregated network of the TARGET dataset. **A** to **G** show scatter plots by community**. H)** show the scatter plot of the complete network (6,032 edges).

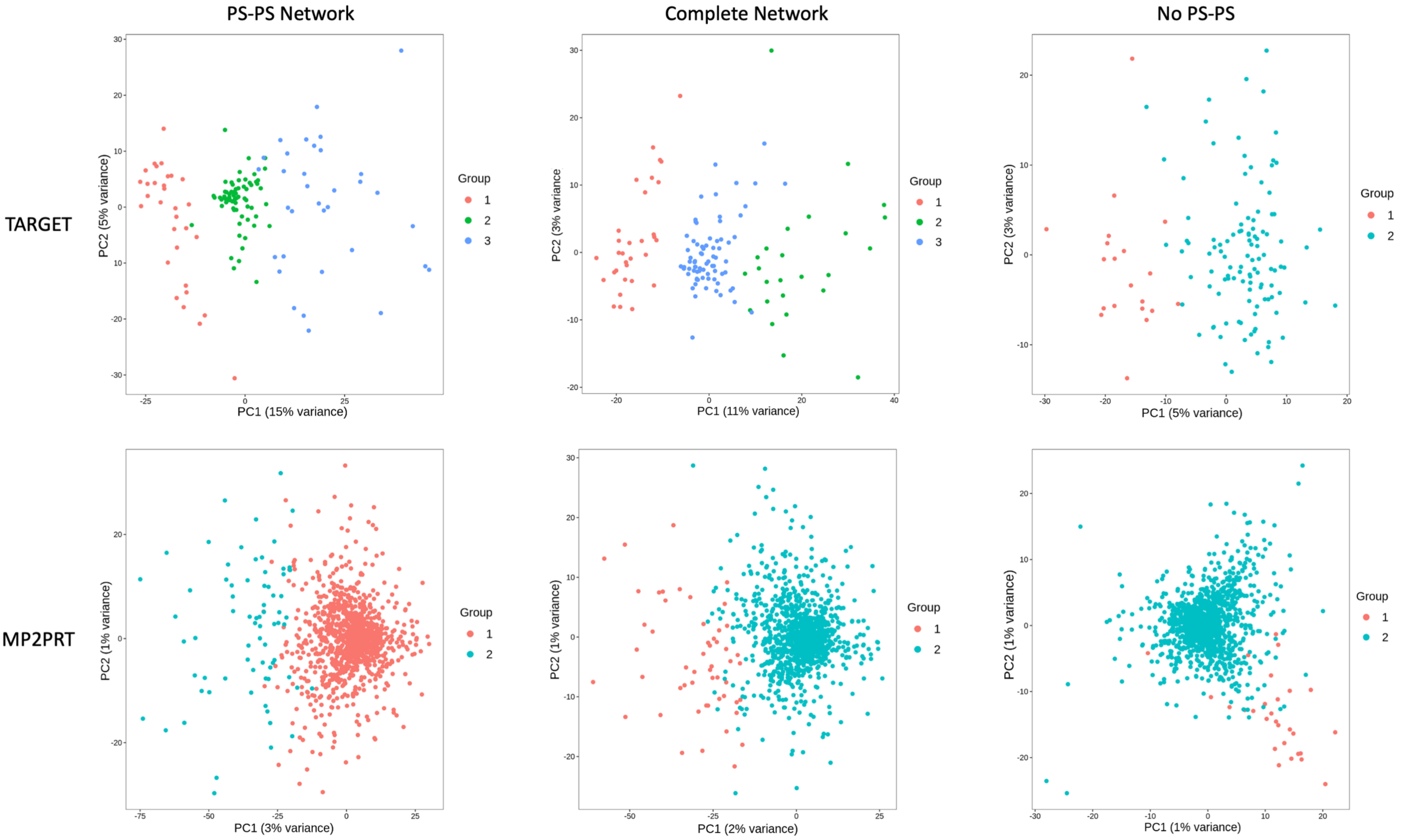


**Fig. S3.** Principal components analysis performed on different network subsets. Top figures show the analysis on the TARGET network, bottom figures show the analysis using the MP2PRT data.

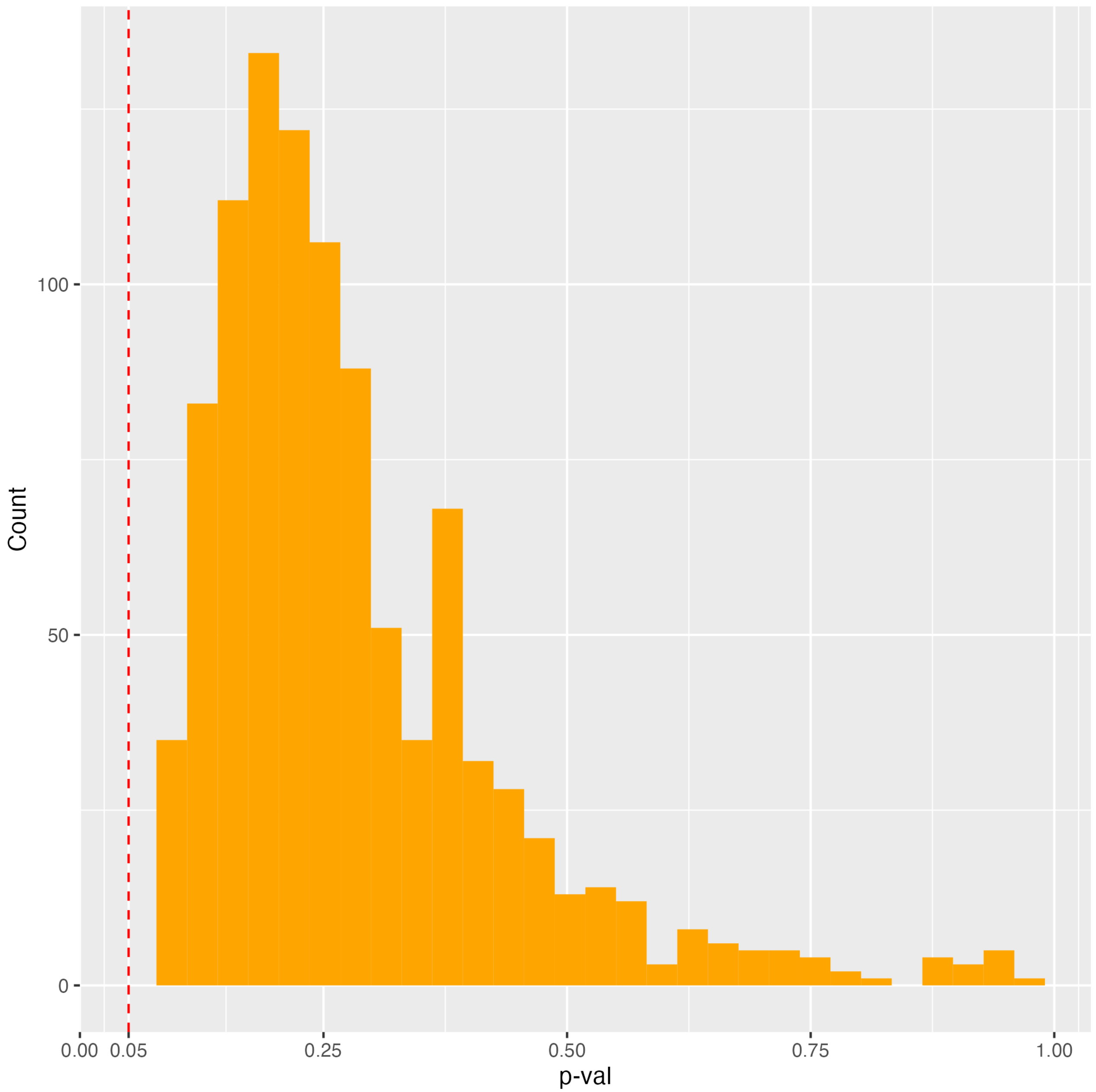


**Fig. S4.** Histogram of p-values from Kaplan-Meier analysis of clusters formed by randomly subsetting (1,000 times) 1,508 edges from the complete TARGET network.

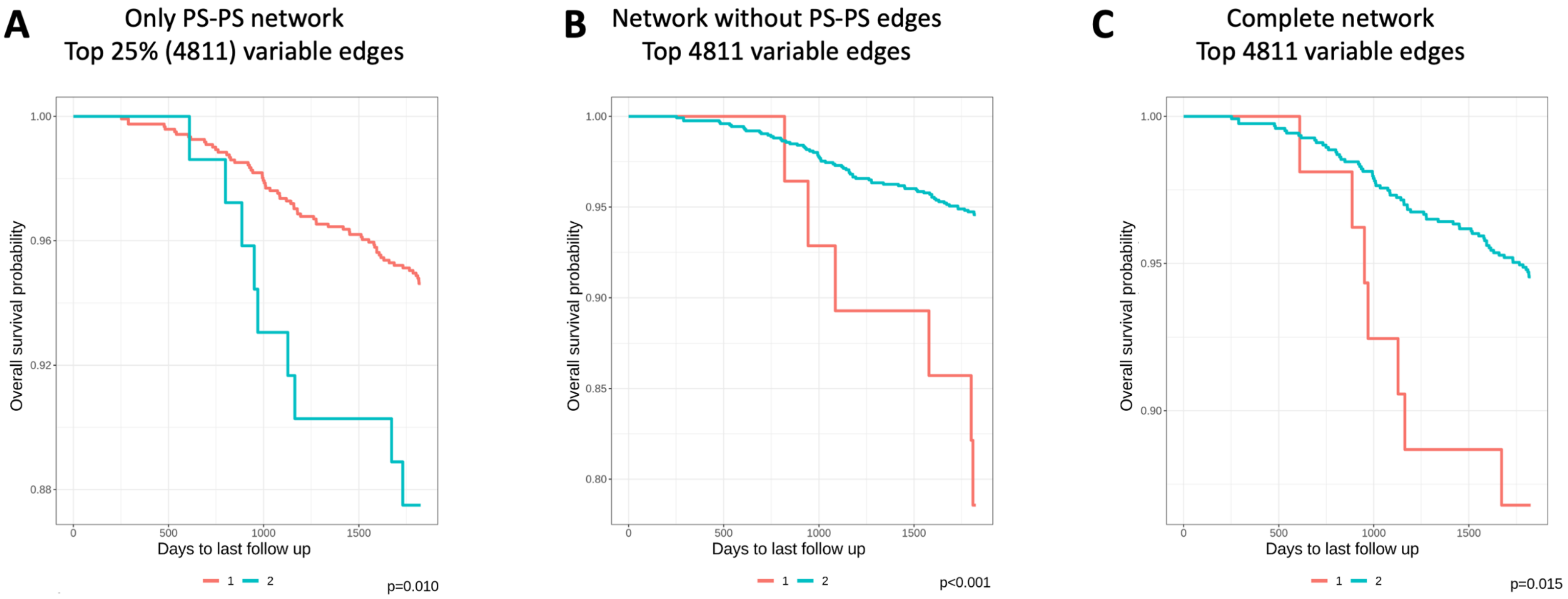


**A**

| | | | | |
|---|---|---|---|---|
| **1** | | | | |
| At Risk | 1212 | 1207 | 1187 | 1159 |
| Events | 0 | 5 | 25 | 46 |
| **2** | | | | |
| At Risk | 72 | 72 | 67 | 65 |
| Events | 0 | 0 | 5 | 7 |

**B**

| | | | | |
|---|---|---|---|---|
| **1** | | | | |
| At Risk | 28 | 28 | 26 | 25 |
| Events | 0 | 0 | 2 | 3 |
| **2** | | | | |
| At Risk | 1256 | 1251 | 1228 | 1199 |
| Events | 0 | 5 | 28 | 50 |

**C**

| | | | | |
|---|---|---|---|---|
| **1** | | | | |
| At Risk | 53 | 53 | 49 | 47 |
| Events | 0 | 0 | 4 | 6 |
| **2** | | | | |
| At Risk | 1231 | 1226 | 1205 | 1177 |
| Events | 0 | 5 | 26 | 47 |

**Fig. S5.** Kaplan-Meier analysis of clusters in MP2PRT data. **A)** Top 25% (4,811 edges) most variable PS-PS edges **B)** Top 4,811 most variable edges from the network without PS-PS edges. **C)** Top 4,811 most variable edges from the complete network containing all classes of edges.

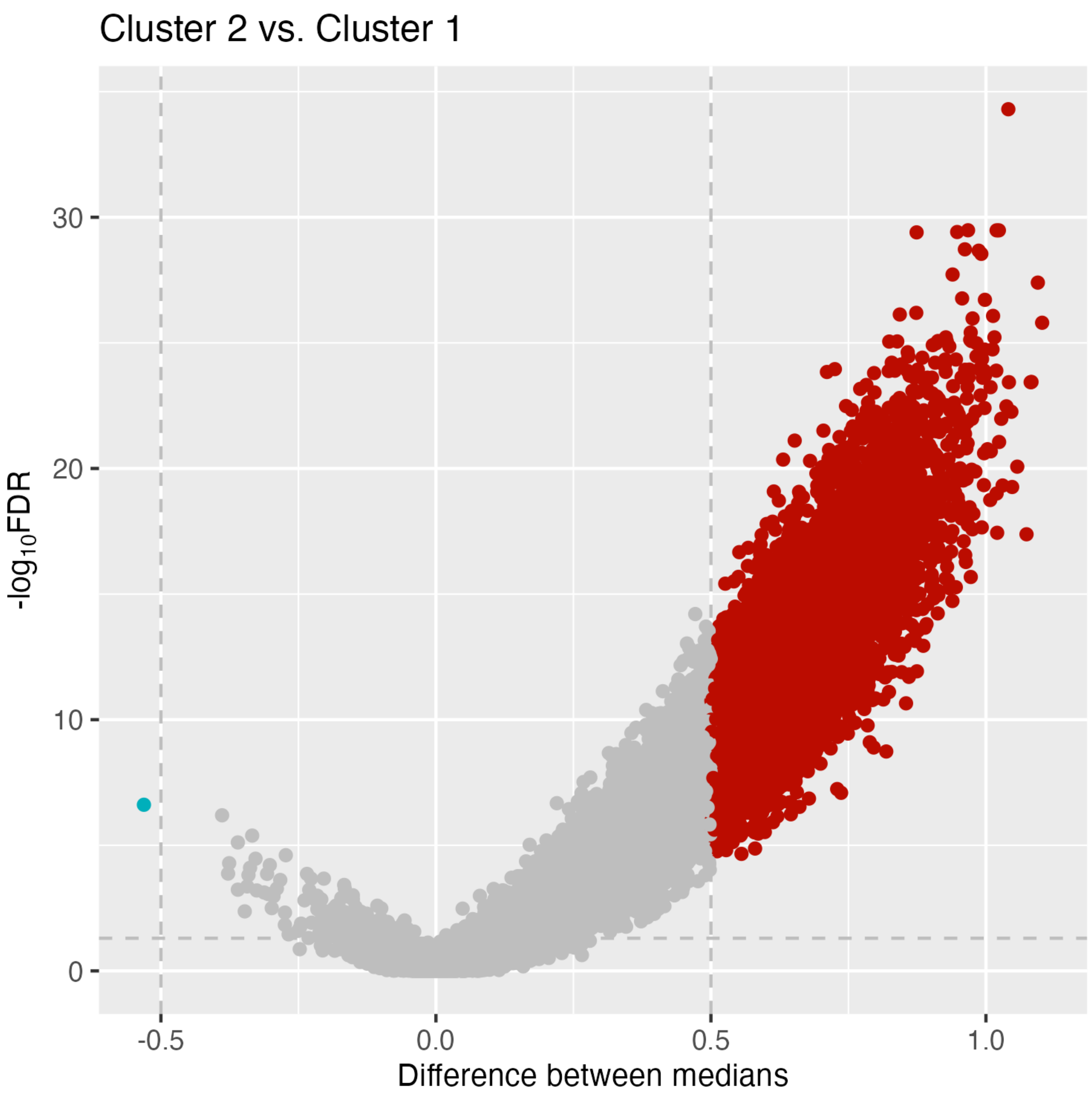


**Fig. S6.** Volcano plot showing the differential co-expression analysis between clusters in the MP2PRT data formed by the analysis of the $PG_{nets}$.

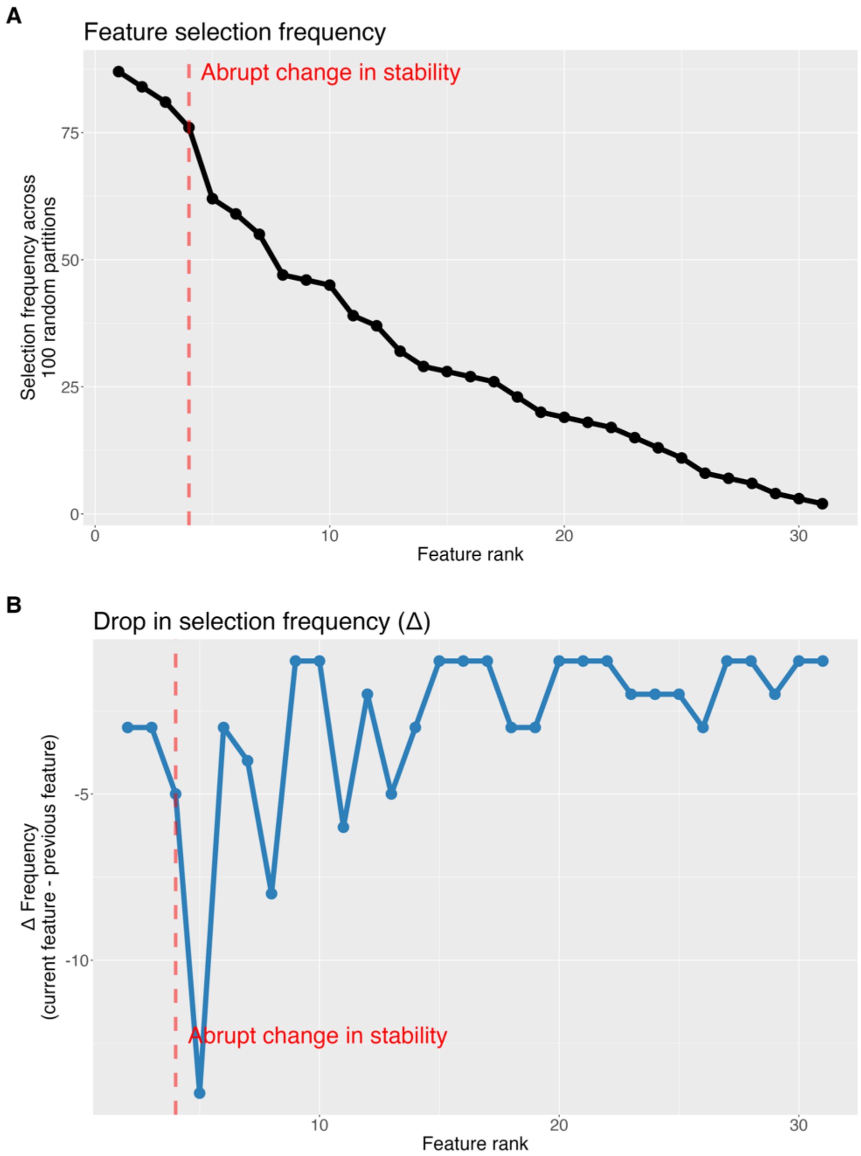


**Fig. S7.** Feature stability across random partitions. **(A)** Features were ranked by how frequently they were selected across 100 LASSO-based feature selection iterations using 50%-sampled random partitions. A steep drop in selection frequency after the top three features indicates a transition from highly stable to less consistently selected features. **(B)** First-order differences (Δ) in selection frequency between consecutive features show a sudden and marked drop at the same point, reinforcing the existence of an inflection point in feature stability. The dashed red line and annotation highlight this abrupt change, supporting the decision to retain only the top three most stable features for downstream modeling.

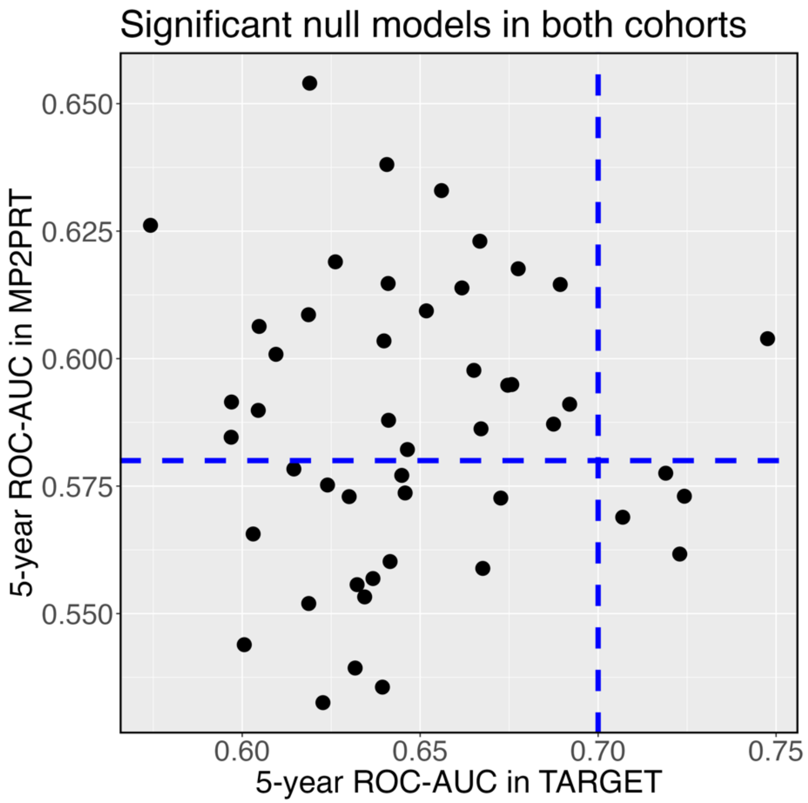


**Fig. S8.** 5-year ROC-AUC values in TARGET and MP2PRT for the 48 null models that consistently achieved significant risk stratification in both cohorts. Each point represents a null model. Dashed lines indicate the 5-year ROC-AUC values obtained by the real model in each cohort. Only one null model exceeds the performance of the real model in both datasets.

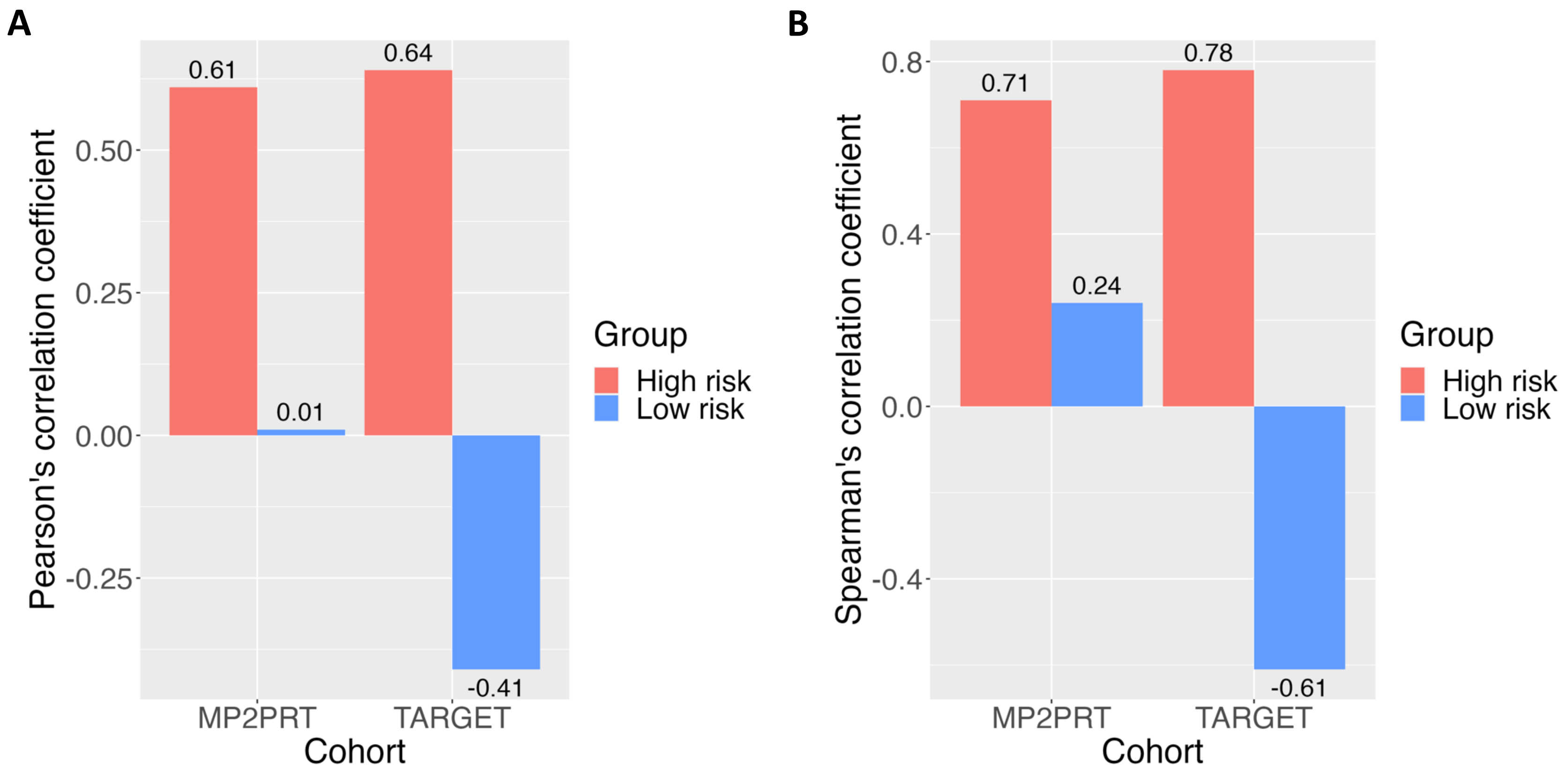


**Fig. S9.** Coefficients of Pearson (**A**) and Spearman (**B**) correlations between samples of high and low survival risk by stratification using the single sample edge weights of *RPL7P10-RPS3AP36*.
